# Coupling microdroplet-based sample preparation, multiplexed isobaric labeling, and nanoflow peptide fractionation for deep proteome profiling of tissue microenvironment

**DOI:** 10.1101/2023.03.13.531822

**Authors:** Marija Veličković, Thomas L. Fillmore, Kwame Attah, Camilo Posso, James C. Pino, Rui Zhao, Sarah M. Williams, Dušan Veličković, Jon M. Jacobs, Kristin E. Burnum-Johnson, Ying Zhu, Paul D. Piehowski

## Abstract

There is increasing interest in developing in-depth proteomic approaches for mapping tissue heterogeneity at a cell-type-specific level to better understand and predict the function of complex biological systems, such as human organs. Existing spatially resolved proteomics technologies cannot provide deep proteome coverages due to limited sensitivity and poor sample recovery. Herein, we seamlessly combined laser capture microdissection with a low-volume sample processing technology that includes a microfluidic device named microPOTS (Microdroplet Processing in One pot for Trace Samples), the multiplexed isobaric labelling, and a nanoflow peptide fractionation approach. The integrated workflow allowed to maximize proteome coverage of laser-isolated tissue samples containing nanogram proteins. We demonstrated the deep spatial proteomics can quantify more than 5,000 unique proteins from a small-sized human pancreatic tissue pixel (∼60,000 µm2) and reveal unique islet microenvironments.

## INTRODUCTION

Biological tissues contain a wide variety of cell types and unique microenvironments with distinct functions^1, 2^. Widely used bulk analysis approaches average out the biological process across whole cell populations, and as such, obscure cellular heterogeneity and spatial localization^3^. Thus, retaining the cellular heterogeneity with high spatial resolution is of great interest in biomedical research. Single-cell and spatial proteomic technologies are becoming increasingly important to understand and discern cellular heterogeneity^3^.

Recently, increasing attention has been focused to enable spatial profiling of low-input samples, which became another challenge for obtaining deep proteome coverage considering the sample loss during sample preparation. Although the various technique utilizes different processing methods^4-8^, the common goal to all of them is a miniaturization of sample preparation to minimize samples loss and maximize digestion kinetics. One promising technique that demonstrated the ability to effectively analyze bottom-up proteomic samples with limited numbers of cells and even down to a single mammalian cell^1, 9-12^, as well as a top-down analyses of a small number of mammalian cells ^13^ is a Nanodroplet Processing in One pot for Trace Samples (nanoPOTS). Even though the nanodroplet processing platform illustrates the power of sample processing miniaturization, it utilizes technology that is not easily accessible for wide research community usage. To overcome that limitation, we scaled up our microfluidic nanoliter operation platform to microdroplet volume by designing larger sized microwells, which can accommodate tissue sample processing from near single-cell to multicell levels in up to 2 µL processing volume. By developing a microfluidic device that operates in low-microliter range and utilizes commercially available laboratory micropipettes, we overcame barriers to adoption that we faced with the nanoPOTS platform, such as the demand for highly customized nanoliter pipetting platform and highly skilled personnel to operate the nanoPOTS platform^14, 15^. Previously, microPOTS technology was utilized for the analysis of ∼25 cultured HeLa cells and ∼10 cells from mouse liver thin sections, where ∼1,800 and 1,200 unique proteins were identified, respectively, with high reproducibility ^15^. This technology was also employed to identify proteomic changes in ∼200 Barrett’s esophageal cells, where >1,500 protein groups were confidently identified, achieving a high reproducibility with a Pearson’s correlation coefficient of R > 0.9 ^14^.

Another limitation of low abundant protein profiling is lack of required sensitivity for MS-based measurements. Proteins are typically identified using a data-dependent acquisition (DDA) label-free approach, which is biased towards selecting peptides with the strongest signal for fragmentation^16^, hence the low-abundant protein quantification is unreliable due to a small number of MS/MS spectra being identified^17^. Although the label-free approach has shown promising results^5, 10, 18-22^, recent advances in sample processing and analysis have mitigated major limitations in MS detection sensitivity^9, 23-30^. Tandem mass tag (TMT) isobaric labeling became a widely used sample multiplexing approach designed to enable identification and precise quantitation of peptides. Several variations on the standard TMT approach have been developed to improve detection of low abundant proteins^9, 25^ and post-translational modifications^23^. Common to all those approaches, is the addition of a carrier channel to the multiplex, which contains a mixture of cells or tissues that mimics the experimental samples with ratio (ratio between the carrier channel and the sample channel) of 30–500-fold.^9^ Another approach that was recently developed that enables a dramatic increase in proteome coverage and enhancement of MS detection sensitivity, is a multidimensional LC separation that includes nanoflow fractionation with an online concatenation process. Previously, a coverage of > 6,000 proteins has been obtained from ∼650 HeLa cells and 10 single human pancreatic islets (∼1,000 cells) utilizing a nanowell-mediated two-dimensional (2D) LC approach^30^. Our in-house built nanoflow Fractionation and Automated Concatenation (nanoFAC) 2D LC platform enabled in-depth proteomic analysis of small-sized samples allowing comprehensive proteome characterization of >6,600 proteins with only 100-ng HeLa digest, equivalent to ∼500 HeLa cells^29^.

To address the difficulty of measuring low-amount protein levels at the spatial scale, we advanced our microPOTS technology by integrating the TMT carrier channel concept into the sample prep workflow, then combined it with our in-house built nanoflow fractionation system to further improve the overall proteome coverages. This effort resulted in a unique platform that can provide deep proteomic profiling of micron-scale tissue samples. Successful implementation of our advanced microdroplet processing technology resulted in quantification of 53,710 unique peptide sequences that mapped to >5,000 unique proteins from a 125,000 µm^2^ rat pancreas tissue voxel. Subsequently, we employed the same technology to image a specific tissue region of a human pancreatic tissue section to gain biological insights on the pancreatic islet microenvironment. Using the optimized microdroplet processing protocol with the TMT carrier concept, we were able to dramatically increase the sensitivity and depth of proteome coverage. Hence, from 60,000 µm^2^ laser captured micro-dissected human pancreas tissue section, we identified 52,000 unique peptide sequences that map to >5,500 unique proteins.

### Experimental section

#### Reagents and Chemicals

Polypropylene microwell chips with 2.2 mm wells diameter were manufactured on polypropylene substrates by Protolabs (Maple Plain, MN). LC-MS grade water, formic acid (FA), iodoacetamide (IAA), Triethylammonium bicarbonate (TEAB), TMT-10plex and TMT11-131C reagents, Anhydrous acetonitrile, Tris(2-carboxyethyl)phosphine hydrochloride (TCEP-HCl), and 50% Hydroxylamine (HA) were purchased from Thermo Fisher Scientific (Waltham, MA). N-Dodecyl β-d-maltose (DDM), DMSO (HPLC grade), Phosphate-Buffered Saline (PBS), and PAS staining kit were purchased from Sigma-Aldrich (St. Louis, MO). Both Lys-C and trypsin were purchased from Promega (Madison, WI). Ethanol was purchased from Decon Labs, Inc (King of Prussia, PA).

#### Tissue collection

To evaluate and demonstrate utility of our enhanced microPOTS technology, we used rodent and human models of pancreas tissue. Sprague Dawley rats (Crl:CD[SD]) were obtained from Charles River Laboratories and were 7-8 weeks of age when pancreas was collected. Animals were euthanized by exsanguination while under isoflurane gas anesthesia, and the pancreas was collected immediately and snap-frozen in optimal cutting temperature compound (OCT). For the alkylation optimization experiments we used rat gastrocnemius muscle tissue from Sprague Dawley rats housed at PNNL according to NIH and institutional guidelines for the use of laboratory animals. Dissected tissues were immediately plunge frozen in LN2 and stored at –80 °C until use. All protocols for this study were reviewed and approved by the Institutional Animal Care and Use Committee of Battelle, Pacific Northwest Division. Human pancreas tissue that was used for microPOTS imaging application was obtained from a 17-year-old male donor. Donor was selected based on our eligibility criterion (dx.doi.org/10.17504/protocols.io.yxmvmnye5g3p/v2). Organ recovery and tissue processing were performed per our standard protocol (dx.doi.org/10.17504/protocols.io.n2bvj6dnblk5/v1). Briefly, pancreas was sliced into 0.5cm-thick tissue segments, subdivided, and immediately frozen in Carboxymethylcellulose (CMC, prepared per dx.doi.org/10.17504/protocols.io.br4fm8tn) in Cryotray molds that were prechilled on dry ice/isopentane slurry. Frozen CMC tissue blocks were stored at -80°C until sectioning.

#### Cryosectioning

OCT embedded rat pancreas tissue was cryosectioned to twelve-micrometer-thick sections using cryostat and thaw-mounted onto polyethylene naphthalate (PEN) membrane slides. The OCT was removed from the tissue sections by immersing slides into 50% ethanol for 30 seconds followed by immersion into the gradient of ethanol solutions (70%, 96%, and 100% ethanol) for 30 seconds each change, to fix the tissue sections.

CMC embedded human pancreas tissue was cut to ten-micrometer-thick slices using cryostat and collected on PEN membrane slides. Samples were washed with the gradient of ethanol solutions (70%, 96%, and 100% ethanol, respectively) for 30 seconds each change, to dehydrate the tissue sections and to remove embedding material.

#### Laser capture microdissection (LCM)

Sample dissection and collection were completed using a PALM Technologies (Carl Zeiss MicroImaging, Munich, Germany) which contains a PALM MicroBeam and RoboStage designed for high-precision laser micromanipulation in micrometer range and a PALM RoboMover that collects dissected samples directly into microwells of the microPOTS chip. For sample collection, microwells were preloaded with 3 µl of dimethyl sulfoxide (DMSO) that served as a capturing medium for excised tissue sections.

First, we used the rat pancreas tissue sections to assess our platform for deep spatial proteomics profiling in terms of robustness and reproducibility. We collected five replicates of each acinar and islet tissue, with a tissue area of 125,000 µm^2^ per replicate. That translates to approximately 4-5 voxels of the whole islets collected per replicate and one acinar tissue voxel per replicate. In our experimental design, we also included a carrier sample of roughly equally distributed islet and acinar tissue containing 16-fold more tissue material than in individual samples, which is approximately 2,000,000 µm^2^ of the total tissue area collected.

For the microPOTS imaging experiment, we first stained a 10 µm thick human pancreas section using Periodic Acid-Schiff (PAS) staining kit, following the manufacturer’s protocol. Informed by the islet localizations from the serial PAS-stained section, a 3×3 grid was used to collect an islet and surrounding acinar tissue. The grid was positioned to capture the whole islet in one 200 µm × 300 µm pixel and the surrounding acinar tissue in the other 8 pixels. Voxels were dissected in the grid mode and collected into corresponding microwells of the chip. One carrier sample was also obtained, containing a similar-sized islet and surrounding acinar tissue with a total area equivalent to the entire grid size, which was 540,000 µm^2^.

#### Proteomics sample processing in a microdroplet

The whole sample prep workflow including the subsequent TMT labeling was carried out on-chip by manual pipetting protocol. Evaporation was minimized through the combination of chip cooling during dispensing of reagent solutions and using a humidified chamber for the incubation steps, where the chip was previously sealed with a contactless cover and wrapped in aluminum foil.

Rat pancreas tissue voxels collected in the microPOTS chip were incubated at 75°C for an hour to dry out remaining DMSO solvent. After DMSO was completely evaporated, 2 µl of extraction buffer containing 0.1% DDM, 0.5×PBS, 38 mM TEAB, and 1 mM TCEP was dispensed to each well of the chip, following incubation at 75°C for an hour. Next, 0.5 µl of IAA solution (25 mM IAA in 100 mM TEAB) was added to the samples, and the samples were allowed to incubate at room temperature for 30 min. All samples were subsequently digested by dispensing 0.5 µl of an enzyme mixture (10 ng of Lys-C and 40 ng of trypsin in 100 mM TEAB) and incubated at 37°C for 10 h. TMT-10 and TMT-11 isobaric mass tag reagents were resuspended in anhydrous acetonitrile to a concentration of 6.4 μg/μL. 1 µl of each TMT tag was dispensed to the corresponding well with the sample. The peptide–TMT mixtures were incubated for 1 h at room temperature, and the labeling reaction was quenched by adding 1µl of 5% HA in 100mM TEAB and incubated for 15 min at room temperature. Next, all samples were pooled together into one Eppendorf tube, brought up to the final 1% FA, then centrifuged at 10,000 rpm for 5 min at 25 °C to spin down precipitate. The supernatant was then transferred to an autosampler vial, dried, and stored in -20°C.

Human pancreas tissue samples, collected for the proteomics imaging experiment, were processed following the aforementioned protocol with a slight modification of the alkylation step and TMT labeling strategy. Informed by the overalkylation of the previously processed rat pancreas samples, we used 0.5 µl of 10 mM IAA solution in 100 mM TEAB to reach a final concentration of 2 mM IAA in the reaction. Following our experimental design, peptides were labeled using a TMT 11-plex, leaving the 130N channel empty and using the 131N channel for the carrier sample, 128N channel was used for the islet voxel and the other 8 channels for the acinar tissue voxels.

#### Nanoflow LC-fractionation

Sample was resuspended in 62 μl of 0.1% formic acid in water. The first-dimension high pH fractionation was performed off-line by loading 50 μl of the sample onto a precolumn (150 µm i.d., 5 cm length) using 0.1% formic acid in water at a flow rate of 9 µL/min for 9 min. The sample was then transferred on an LC column (75 µm i.d., 60-cm length). Precolumn and column were packed inhouse with 5-µm and 3-µm Jupiter C18 packing material (300-Å pore size) (Phenomenex, Terrence, USA), respectively. An Ultimate 3000 RSLCnano system (Thermo Scientific) was used to deliver gradient flow to the LC column at a nanoflow rate of 300 nl/min. 10 mM ammonium formate (pH 9.5) in water was used as mobile phase A and acetonitrile as mobile phase B. The peptides were eluted using the following increasing gradients: from 1% to 8% of mobile phase B in 11 min and then to 12% mobile phase B in 18 min, followed by an increase to 30% mobile phase B in 55 min, then to 45% mobile phase B in 22 min, to 65% mobile phase B in 6 min and finally to 95% mobile phase B in 5 min. Peptides eluted from the high-pH nanoLC separation were fractionated using a HTX PAL collect system into the vials preloaded with 25 μl 0.1% formic acid in water, containing 0.01% (m/v) DDM. The PAL autosampler allowed us to automatically perform the concatenation by robotically moving the dispensing capillary among 12 collection vials, hence a total of 96 fractions were concatenated into 12 fractions. Vials were stored at −20 °C until the following low-pH LC-MS/MS analysis.

#### LC-MS/MS peptide analyses

Fractionated samples were separated in the second dimension, using the Ultimate 3000 RSLCnano system (Thermo Scientific), by injected fully into a 20 µL loop and loaded onto a precolumn using 0.1% formic acid in water at a flow rate of 7 µL/min for 5 min. 3 cm-long Jupiter trapping column with 150 µm i.d was inhouse packed using 5 µm C18 packing material (300-Å pore size, Phenomenex, Terrence, USA). Sample was then transferred on an LC column (1.7 µm Waters BEH 130 75 µm idx 30 cm) that was heated at 45°C. Chromatographic separation was performed at 200 nL/min using the following gradient: 1-8% (2.3-12.6 min), 8-25% (12.6-107 min), 25-35% (107.6-117.6), 35-75% (117-122.6 min), and 75-95% (122.6-125.9 min) of Buffer B (0.1% formic acid in acetonitrile). Separated peptides were introduced to the ion source of an Q Exactive HF-X (Thermo Scientific) instrument, where high voltage (2200V) was applied to generate electrospray and ionize peptides. The ion transfer tube was heated to 300 °C and the S-Lens RF level was set to 40. Similar acquision parameters were used for analyzes of the two different pancreas tissue models. Full MS scan of rat pancreas tissue sample fractions were acquired across scan range of 300 to 1,800 m/z at a resolution of 60,000, combined with a maximum injection time (IT) of 50 ms and automatic gain control (AGC) target value of 3e6. Sixteen data-dependent MS/MS scans were recorded per MS scan, at a resolving power of 60,000 combined with a maximum injection time of 120 ms and AGC target value of 2e5, with an isolation window of 0.7 m/z. A dynamic exclusion time was set to 45 s to reduce repeated selection of precursor ions. For the human pancreas tissue fractions, precursor ions from 300 to 1,800 m/z were scanned with a mass resolution of 60,000 combined with an IT of 20 ms and (AGC) target of 3e6. The MS/MS spectra were acquired in data-dependent mode where twelve most intense precursor ions were fragmented and recorder at a resolving power of 45,000 combined with a maximum injection time of 100 ms and AGC target value of 1e5, with an isolation window of 0.7 m/z. A dynamic exclusion of 45 s was used.

#### Data Analysis

Thermo RAW files were processed using mzRefinery to correct for mass calibration errors^31^. Spectra were then searched with MS-GF + v9881^32, 33^ to match against the Uniprot human database downloaded in March 2021 (20,371 proteins), combined with common contaminants (e.g., trypsin, keratin). A partially tryptic search was used with a ± 20 parts per million (ppm) parent ion mass tolerance. TMT global proteomics data processing was performed as we previously described. ^34^

Gene set enrichment analysis was performed on rat pancreas data using the gseGO^35^ function of the ClusterProfiler R package in conjunction with genome wide annotations for rat (available from org.Rn.eg.db), with the Gene Ontology Biological Process^36, 37^ as the reference database. To obtain the ranked list, we used the MSnSet.utils and limma R^38^ package to compute a t-test comparison between Acinar and Islet samples, then filtered the original 5021 genes down to those with BH adjusted p-values below 0.05, leaving 2339 differentially expressed genes. Finally, we ranked the differentially expressed genes according to the log-fold change of Islet minus Acinar, in descending order, to obtain the input for GSEA.

Human pancreas data was visualized using a python class designed for proteomics imaging data. The datasets are available from Synapse with Synapse ID syn30389393.

## Results and discussion

### MicroPOTS platform for deep spatial proteomics

Figure 1 depicts the workflow for our deep spatial proteomics platform, which combines the LCM-based high resolution sample isolation with the microPOTS sample preparation approach for high sample recovery, a TMT carrier channel strategy, and nanoscale fractionation system with automated concatenation, to dramatically increase the depth of proteome coverage. This approach combines in-depth quantitative information on protein abundance with spatial context for untargeted investigations of biological tissues. The coupling of the microPOTS chip with LCM provides simple, robust and efficient voxel collection. Using the chip with larger well diameters, such as microwell chips with 2.2 mm well diameter, ensures a higher collection success rate, compared to 1.2 mm nanowell collection. Similar to nanoPOTS chip, the microPOTS chip allows the direct visualization of the sample with the LCM microscope to confirm sample capture (see Supporting Information: Figure 1). In terms of the sample processing, the low µL reaction volume in microPOTS chip improved kinetics, reduced contaminants and side-reactions. The microPOTS platform retains some advantages of the nanoPOTS platform, such as high digestion efficiency and minimal sample losses during sample preparation. As opposed to our nanodroplet-based proteomic approach, which requires highly customized sample processing equipment, our microdroplet-based proteomic approach operates in the low-microliter range hence the samples can be processed manually using standard pipettes. As such, microPOTS is a low-cost technology, designed to be easily adopted by other research labs. Next, we utilized a multiplexing strategy by combining TMT isobaric labeling and a TMT carrier channel with the microPOTS technology to improve proteome depth. Collection and labeling of large enough samples enabled us to utilize our custom-built nano-scale fractionation 2D LC system and dramatically increase the overall sensitivity and coverage of the platform. The resolving power and overall peak capacity dramatically increased by using two-dimensional separation, which in combination with the nanofractionation decreased sample complexity in each of the fractions and therefore enabled us to maximize proteome coverage of small-sized samples.

**Figure 1.**
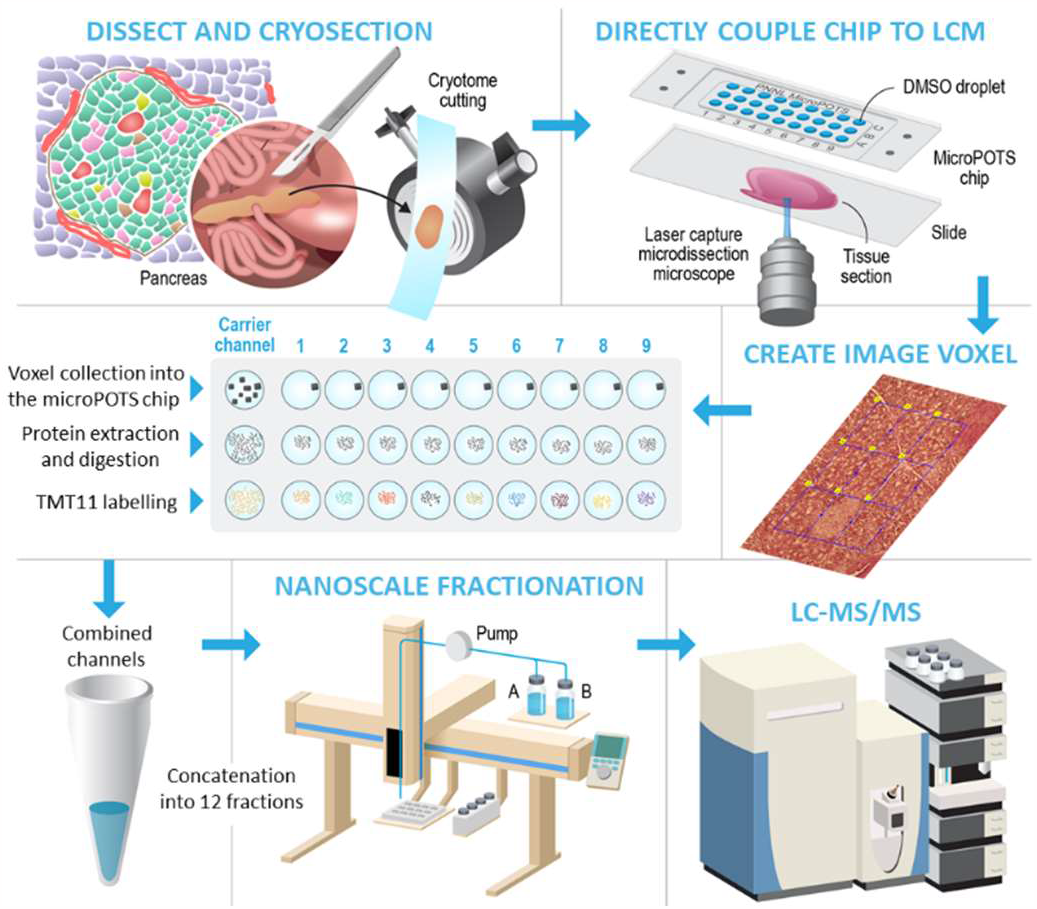
Schematic overview of the deep spatial proteomics analysis pipeline. Our advanced platform for micron-scale proteomics integrates the TMT labeling strategy with carrier channel into the microPOTS sample prep workflow and utilizes a home-built nanoflow fractionation system to maximize proteome coverage of low-input samples.

### Implementation and evaluation of our deep spatial proteomics platform

To demonstrate that our recently developed platform for microscale proteomics can provide in-depth, quantitative analyses of limited sample amounts, we employed it to compare proteome profiles of two distinct pancreas tissue types. Exocrine pancreas tissue (acinar) and endocrine pancreas tissue (islet) were dissected and collected directly into a microPOTS chip following the experimental design detailed in the Experimental Section. To assess the robustness of our platform, we first looked at various analytical figures of merit. We found that our microPOTS sample processing approach provided great efficiency with a missed cleavage rate <15%. TMT labeling efficiency was evaluated by comparing the number of unlabeled peptides with the labeled peptides^39^, and the labeling efficiency was >99%, which indicated that our microPOTS platform provides as highly efficient TMT labeling as the nanoPOTS platform. Looking across the dataset, the sample channels showed between 15 and 26-fold median abundance difference when compared to the carrier channel loaded with a 16-fold excess sample by area (fig. 2-a). Looking more closely at the sample channels, we can see that median intensities vary (with mean 18444.47 and standard deviation 3444.564) indicating peptide yields are comparable when loading equal tissue area (fig. 2-b). Additionally, we looked at missing data at the peptide and protein levels of the 11-plex set. Among the identified, > 99% of proteins and > 95% peptides were quantified without any missing values in the channels (fig. 2-c), demonstrating good data completeness achievable using TMT with a carrier channel, which is especially important considering that peptide loading amounts were at low-μg level. We also assessed performance of our custom nanofractionation system. We identified 53,710 unique peptide sequences that correspond to 5,202 unique proteins, whereas 78% of unique peptide identifications were found within a single fraction and 95% of peptides were found in two or fewer fractions, indicating high fractionation efficiency (fig. 2d). Peptide were evenly distributed across the fractions, with each fraction yielding between 5,300 to 6,200 peptides, showing low fraction-to-fraction variation (fig. 2e). Further, we computed the coefficient of variation (CV) for 5 islet and 5 acinar tissue replicates at the protein level (fig. 2-f), and the results showed that variation among the islet replicates was comparatively small, median CV <15%, indicating good reproducibility of the platform. We found that the CV for the acinar replicates, median CV 27%, was much higher relative to the islet. One potential reason for the different grouping of the acinar replicates and their high CV is that samples were collected from different tissue locations, indicating that the acinar tissue is not as homogeneous as we expected it to be, whereas islet samples were a pool of islets creating a more homogenous sample.

**Figure 2.**
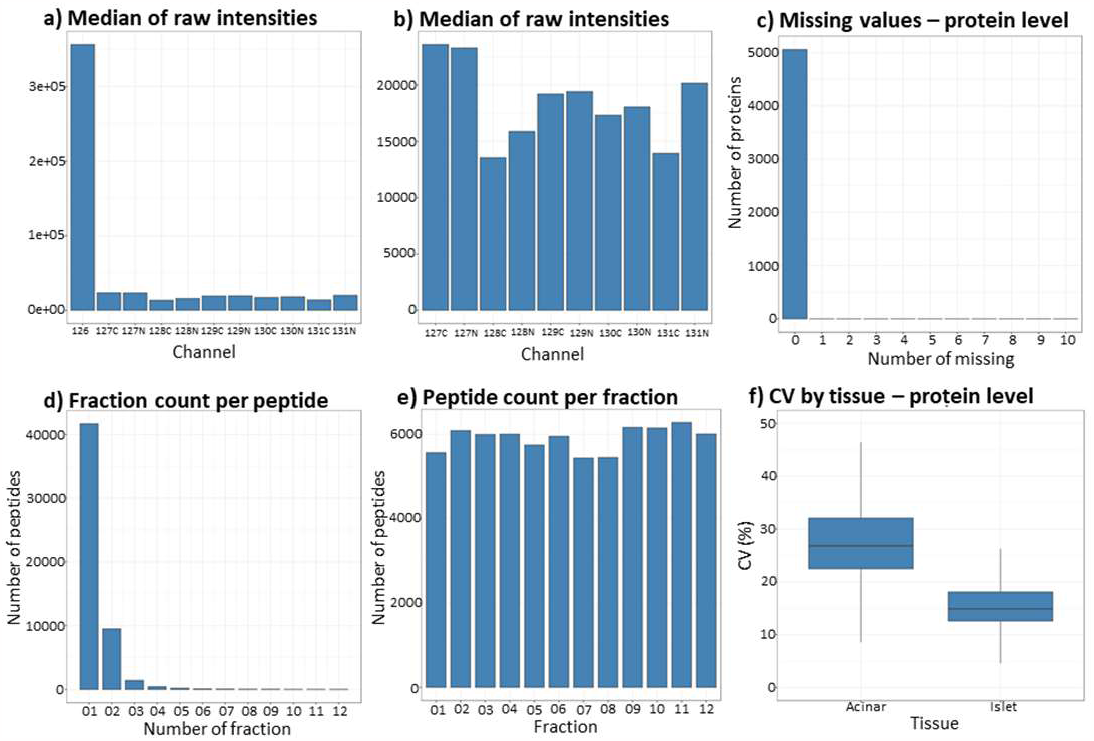
Evaluation of the deep spatial proteomics platform. a-b) The bar graphs show median relative abundance in the 11-plex set. Carrier channel loaded with a 16-fold excess sampling area was labelled with the TMT 126. c) Missing data at the protein levels. d) The fraction count in which a given peptide was detected. e) Sampling depth (peptide count) for each fractionation. f) Median CVs calculated across 5 replicates of each of two pancreas functional units.

After assessing the microPOTS figures of merit, we compared proteomic profiles generated to evaluate the platform’s ability to accurately identify biological differences. A principal component analysis (PCA) was employed on the data from all 10 samples to visualize whether a clear separation exists between islet and acinar proteomic profiles. This analysis clearly separated islet and acinar tissues forming two separate clusters in the first PC, accounting for 2/3 of the variability in the dataset (fig. 3-a). While five islet replicates clustered closely together, the second PC separated acinar replicates into two groups accounted for 10% of the variation. Next, we used Linear Models for Microarray Data (LIMMA) to compare the two different tissue types. As expected, given the disparate functions of these tissue types, we found 2,339 differentially expressed proteins. We then took the genes with an adjusted p-value below an FDR of 0.05, hierarchical clustering was used to identify differences in gene expression between the islet and acinar tissue types. Generated gene expression heat map showed a distinct cluster of genes for islet tissue data with higher abundances relative to acinar tissue data, indicating different biological functions of the two tissue types (**fig. 3b**). Consistent with known differences in function, islet samples overexpressed genes related to cell-cell signaling and secretion, while acinar tissue was enriched in metabolic terms (**fig. 3c**).

**Figure 3.**
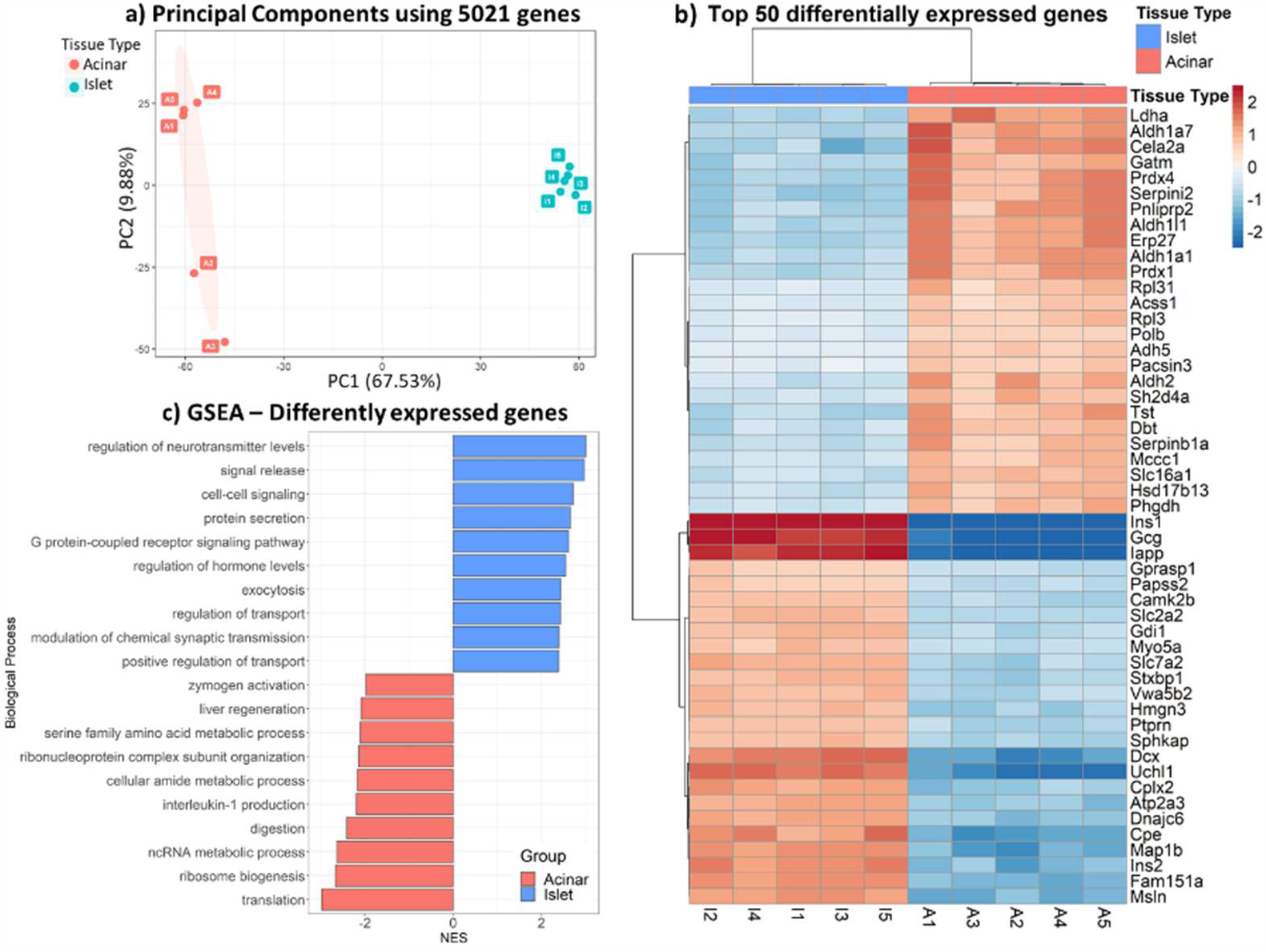
(a) PCA showing sample variation in 2D principal component space for islet and acinar proteomic profiles (b) Heatmap depicts hierarchical clustering of top 50 differently expressed genes in 2 functional units of the pancreas (c) Gene set enrichment analysis for 2339 genes differentially expressed in endocrine and exocrine pancreas.

### In-depth proteome imaging of human islet microenvironment

Having established the quality performance of our microPOTS platform, we sought to demonstrate its utility as a proteome imaging platform. To do this, we captured the entire islet in one pixel and acinar tissue in the other 8 pixels in order to investigate the acinar microenvironment in the region surrounding the islet, figure 4. From 200 µm × 300 µm pancreas tissue, we reliably identified 52,000 unique peptide sequences that map to > 5,500 unique proteins. We then mapped known tissue-specific proteins to see if the localization of these markers aligns with the functional role. As was expected, endocrine hormones insulin and glucagon were predominantly present in the endocrine (islet) region and, on the other hand, digestive enzymes alpha-amylase 2B (AMY2B) and pancreatic triacylglycerol lipase (LIPP) were localized in the acinar region of the exocrine pancreas. Figure 4 shows the log2 abundance values of insulin and glucagon hormones highly expressed in the islet, as well as the presence of acinar tissue markers AMY2B and LIPP, across all 9 imaged tissue pixels. This imaging experiment demonstrates the platform’s ability to provide proteomic profiles at a high spatial resolution, allowing us to image an important region of the pancreas with unprecedented depth of proteomic coverage.

**Figure 4.**
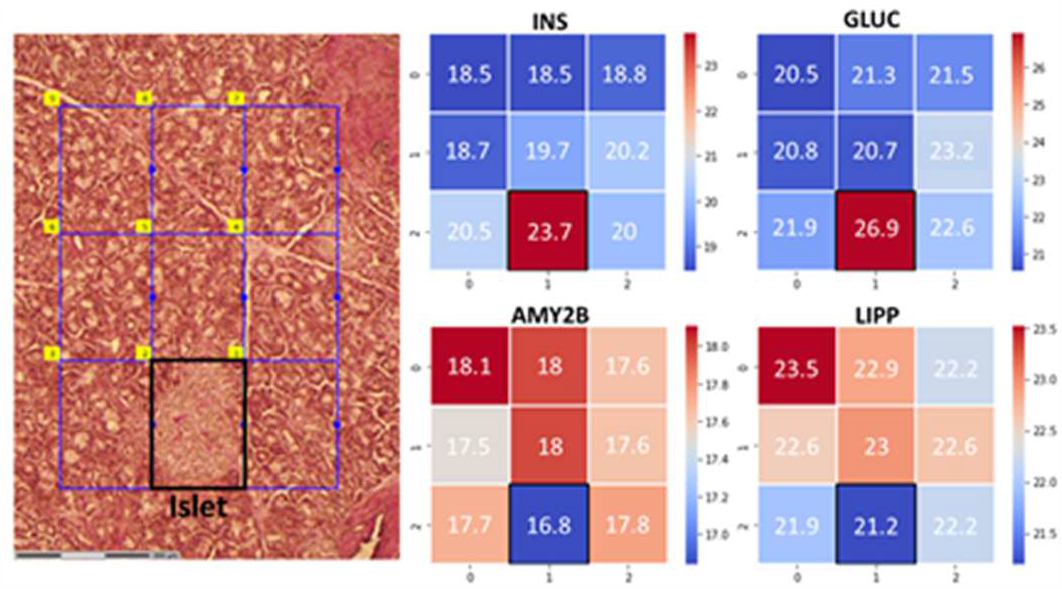
Proteome imaging data obtained using our advanced microPOTS platform for deep spatial proteomics profiling.

### Conclusions

Here we introduce a novel, microdroplet-based platform that combines LCM, microPOTS processing, multiplexing labeling with a carrier channel, and nanoFAC to dramatically increase the sensitivity and depth of proteome coverage for small, spatially resolved, samples. We have demonstrated the performance and reproducibility of our platform by obtaining deep protein coverage and matching functional units of the pancreas with their protein profiles at 200 µm spatial resolution.

Our advanced microPOTS technology can be applied to virtually any other biological sample and hence it can be broadly applied across biomedical research. Microliter sample prep technology can be performed with a micropipette without the requirement of nanoliter liquid handling robot. This, the low-cost technology is easily implanted and adopted by other research labs. Further, the nanoFAC system could be assembled from commercially available instrumentation and parts. Currently, this technology uses a TMT 11-plex study design, future experiments can take advantage of TMT 18-plex reagents to further increase the throughput.

## ACKNOWLEGMENT

This work was supported by the NIH HubMAP initiative grant to JL NIH UH3CA255132 and Pfizer Worldwide Research and Development. Much of this research was performed using the Environmental Molecular Sciences Laboratory, a DOE Office of Science User Facility sponsored by the Office of Biological and Environmental Research and located at Pacific Northwest National Laboratory (PNNL). PNNL is operated for the DOE by Battelle Memorial Institute under contract DE-AC05-76RLO1830.

## Supporting Information

**Supporting figure 1:**
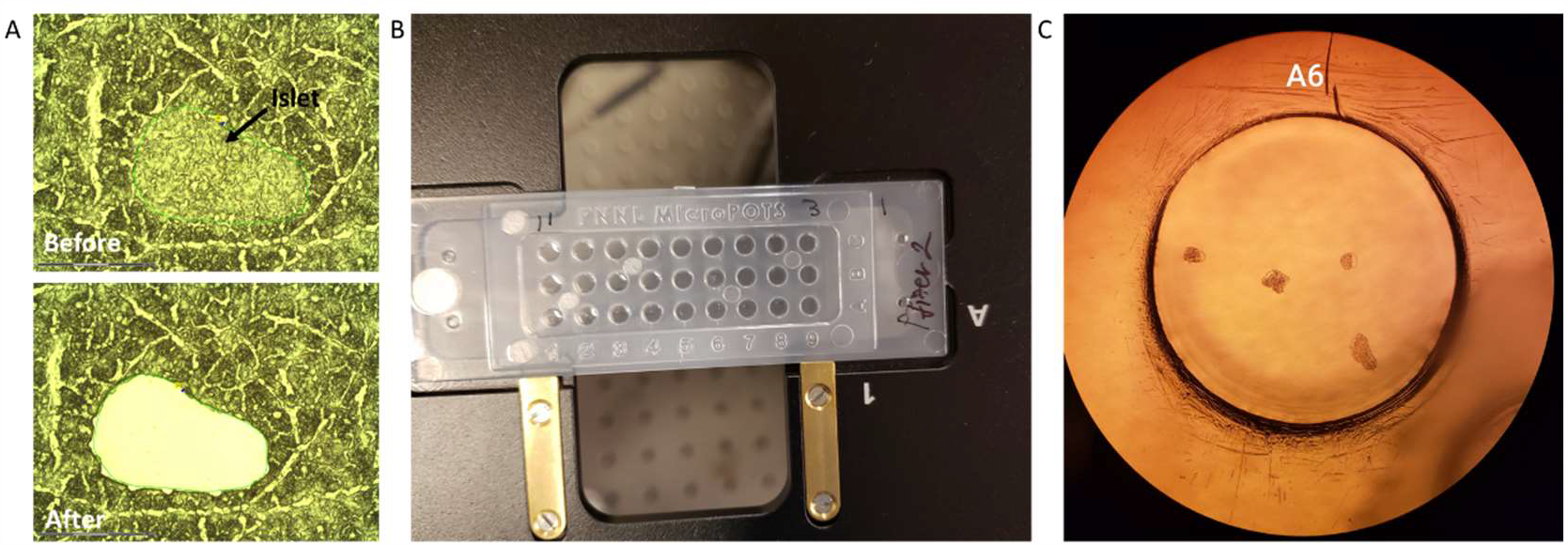
Dissection and collection of the pancreas tissue into the microPOST chip. A) Islet region before and after LCM cut B) Microchip with collected pancreas tissue samples. Microwells were preloaded with DMSO that served as a capturing medium. C) Islet collected into microwell A6, observed under the Zeiss LCM microscope.

## REFERENCES

1. Zhu, Y., et al., Spatially Resolved Proteome Mapping of Laser Capture Microdissected Tissue with Automated Sample Transfer to Nanodroplets. Molecular & Cellular Proteomics, 2018. 17(9): p. 1864–1874.

2. Mao, Y.H., et al., Spatial proteomics for understanding the tissue microenvironment. Analyst, 2021. 146(12): p. 3777–3798.

3. Yang, L.W., J. George, and J. Wang, Deep Profiling of Cellular Heterogeneity by Emerging Single-Cell Proteomic Technologies. Proteomics, 2020. 20(13).

4. Sandbaumhuter, F.A., et al., Well-Plate muFASP for Proteomic Analysis of Single Pancreatic Islets. J Proteome Res, 2022. 21(4): p. 1167–1174.

5. Li, Z.Y., et al., Nanoliter-Scale Oil-Air-Droplet Chip-Based Single Cell Proteomic Analysis. Anal Chem, 2018. 90(8): p. 5430–5438.

6. Sielaff, M., et al., Evaluation of FASP, SP3, and iST Protocols for Proteomic Sample Preparation in the Low Microgram Range. J Proteome Res, 2017. 16(11): p. 4060–4072.

7. Leipert, J. and A. Tholey, Miniaturized sample preparation on a digital microfluidics device for sensitive bottom-up microproteomics of mammalian cells using magnetic beads and mass spectrometry-compatible surfactants. Lab Chip, 2019. 19(20): p. 3490–3498.

8. Gebreyesus, S.T., et al., Streamlined single-cell proteomics by an integrated microfluidic chip and data-independent acquisition mass spectrometry. Nat Commun, 2022. 13(1): p. 37.

9. Dou, M.W., et al., High-Throughput Single Cell Proteomics Enabled by Multiplex Isobaric Labeling in a Nanodroplet Sample Preparation Platform. Analytical Chemistry, 2019. 91(20): p. 13119–13127.

10. Zhu, Y., et al., Nanodroplet processing platform for deep and quantitative proteome profiling of 10-100 mammalian cells. Nature Communications, 2018. 9.

11. Piehowski, P.D., et al., Automated mass spectrometry imaging of over 2000 proteins from tissue sections at 100-mu m spatial resolution. Nature Communications, 2020. 11(1).

12. Williams, S.M., et al., Automated Coupling of Nanodroplet Sample Preparation with Liquid Chromatography-Mass Spectrometry for High-Throughput Single-Cell Proteomics. Analytical Chemistry, 2020. 92(15): p. 10588–10596.

13. Zhou, M.W., et al., Sensitive Top-Down Proteomics Analysis of a Low Number of Mammalian Cells Using a Nanodroplet Sample Processing Platform. Analytical Chemistry, 2020. 92(10): p. 7087–7095.

14. Weke, K., et al., MicroPOTS Analysis of Barrett’s Esophageal Cell Line Models Identifies Proteomic Changes after Physiologic and Radiation Stress. Journal of Proteome Research, 2021. 20(5): p. 2195–2205.

15. Xu, K.R., et al., Benchtop-compatible sample processing workflow for proteome profiling of < 100 mammalian cells. Analytical and Bioanalytical Chemistry, 2019. 411(19): p. 4587–4596.

16. He, B., et al., Label-free absolute protein quantification with data-independent acquisition. Journal of Proteomics, 2019. 200: p. 51–59.

17. Zhou, J.Y., et al., Unraveling pancreatic islet biology by quantitative proteomics. Expert Review of Proteomics, 2011. 8(4): p. 495–504.

18. Shao, X., et al., Integrated Proteome Analysis Device for Fast Single-Cell Protein Profiling. Anal Chem, 2018. 90(23): p. 14003–14010.

19. Brunner, A.D., et al., Ultra-high sensitivity mass spectrometry quantifies single-cell proteome changes upon perturbation. Molecular Systems Biology, 2022. 18(3).

20. Liang, Y., et al., Fully Automated Sample Processing and Analysis Workflow for Low-Input Proteome Profiling. Anal Chem, 2021. 93(3): p. 1658–1666.

21. Zhu, Y., et al., Proteomic Analysis of Single Mammalian Cells Enabled by Microfluidic Nanodroplet Sample Preparation and Ultrasensitive NanoLC-MS. Angewandte Chemie-International Edition, 2018. 57(38): p. 12370–12374.

22. Matzinger, M., et al., Robust and Easy-to-Use One-Pot Workflow for Label-Free Single-Cell Proteomics. Anal Chem, 2023. 95(9): p. 4435–4445.

23. Yi, L., et al., Boosting to Amplify Signal with Isobaric Labeling (BASIL) Strategy for Comprehensive Quantitative Phosphoproteomic Characterization of Small Populations of Cells. Analytical Chemistry, 2019. 91(9): p. 5794–5801.

24. Tsai, C.F., et al., An Improved Boosting to Amplify Signal with Isobaric Labeling (iBASIL) Strategy for Precise Quantitative Single-cell Proteomics. Molecular & Cellular Proteomics, 2020. 19(5): p. 828–838.

25. Budnik, B., et al., SCoPE-MS: mass spectrometry of single mammalian cells quantifies proteome heterogeneity during cell differentiation. Genome Biology, 2018. 19.

26. Ctortecka, C., et al., Quantitative Accuracy and Precision in Multiplexed Single-Cell Proteomics. Analytical Chemistry, 2022. 94(5): p. 2434–2443.

27. Cheung, T.K., et al., Defining the carrier proteome limit for single-cell proteomics. Nature Methods, 2021. 18(1): p. 76-+.

28. Schoof, E.M., et al., Quantitative single-cell proteomics as a tool to characterize cellular hierarchies. Nature Communications, 2021. 12(1).

29. Dou, M.W., et al., Automated Nanoflow Two-Dimensional Reversed-Phase Liquid Chromatography System Enables In-Depth Proteome and Phosphoproteome Profiling of Nanoscale Samples. Analytical Chemistry, 2019. 91(15): p. 9707–9715.

30. Dou, M.W., et al., Nanowell-mediated two-dimensional liquid chromatography enables deep proteome profiling of < 1000 mammalian cells. Chemical Science, 2018. 9(34): p. 6944–6951.

31. Gibbons, B.C., et al., Correcting systematic bias and instrument measurement drift with mzRefinery. Bioinformatics, 2015. 31(23): p. 3838–3840.

32. Kim, S. and P.A. Pevzner, MS-GF plus makes progress towards a universal database search tool for proteomics. Nature Communications, 2014. 5.

33. Kim, S., N. Gupta, and P.A. Pevzner, Spectral probabilities and generating functions of tandem mass spectra: A strike against decoy databases. Journal of Proteome Research, 2008. 7(8): p. 3354–3363.

34. Gosline, S.J.C., et al., Proteomic and phosphoproteomic measurements enhance ability to predict ex vivo drug response in AML. Clinical Proteomics, 2022. 19(1).

35. Wu, T., et al., clusterProfiler 4.0: A universal enrichment tool for interpreting omics data. Innovation (Camb), 2021. 2(3): p. 100141.

36. Ashburner, M., et al., Gene Ontology: tool for the unification of biology. Nature Genetics, 2000. 25(1): p. 25–29.

37. Carbon, S., et al., The Gene Ontology resource: enriching a GOld mine. Nucleic Acids Research, 2021. 49(D1): p. D325–D334.

38. Ritchie, M.E., et al., limma powers differential expression analyses for RNA-sequencing and microarray studies. Nucleic Acids Research, 2015. 43(7).

39. Hutchinson-Bunch, C., et al., Assessment of TMT Labeling Efficiency in Large-Scale Quantitative Proteomics: The Critical Effect of Sample pH. Acs Omega, 2021. 6(19): p. 12660–12666.

